# Reliable Phylogenetic Regressions for Multivariate Comparative Data: Illustration with the MANOVA and Application to the Effect of Diet on Mandible Morphology in Phyllostomid Bats

**DOI:** 10.1101/606491

**Authors:** Julien Clavel, Hélène Morlon

## Abstract

Understanding what shapes species phenotypes over macroevolutionary time scales from comparative data requires the use of reliable phylogenetic regression techniques and associated tests (e.g. phylogenetic Generalized Least Squares, pGLS and phylogenetic analyses of variance and covariance, pANOVA, pANCOVA). While these tools are well established for univariate data, their multivariate counterparts are lagging behind. This is particularly true for high dimensional phenotypic data, such as morphometric data. Here we implement well-needed likelihood-based multivariate pGLS, pMANOVA and pMANCOVA, and use a recently-developed penalized likelihood framework to extend their application to the difficult case when the number of traits *p* approaches or exceeds the number of species *n*. We then focus on the pMANOVA and use intensive simulations to assess the performance of the approach as *p* increases, under various levels of phylogenetic signal and correlations between the traits, phylogenetic structure in the predictors, and under various types of phenotypic differences across species groups. We show that our approach outperforms available alternatives under all circumstances, with a greater power to detect phenotypic differences across species group when they exist, and a low risk to improperly detect inexistent differences. Finally, we provide an empirical illustration of our pMANOVA on a geometric-morphometric dataset describing mandible morphology in phyllostomid bats along with data on their diet preferences. Our approach, implemented in the R package mvMORPH, provides efficient multivariate phylogenetic regression tools for understanding what shapes phenotypic differences across species.

---

Along with other areas of biological sciences, the study of phenotypic traits is experiencing the *omics* revolution: the “phenome”, which describes the various phenotypic characteristics of organisms, is defined by a large number of multivariate anatomical and behavioral traits (Deans et al. 2015). For instance, high resolution morphological datasets are becoming increasingly available across the tree of life thanks to computerized tomography techniques (e.g., Cooney et al. 2017; Cross 2017; Felice and Goswami 2018). These datasets have the potential to dramatically improve our understanding of the factors that influence species phenotypes over macroevolutionary time scales.

While high-dimensional phenotypic datasets are being collected at an ever-increasing scale, the statistical machinery necessary to analyze such datasets is lagging behind. In particular, in order to study the relationship between complex phenotypes and putative explanatory factors from comparative data, or to test for differences in phenotypes across species groups, multivariate regressions accounting for statistical non-independence due to shared ancestry, as well as associated tests, are required (pGLS, pMANOVA and pMANCOVA Felsenstein 1985; Grafen 1989; Martins and Hansen 1997; Rohlf 2001, 2006; Revell 2010; Blomberg et al. 2012; Hansen and Bartoszek 2012). In low-dimensional settings (i.e. when the number of traits *p* is small compared to the number of species *n*), it is possible to test for differences in phenotypes across species groups (i.e. to perform pMANOVAs) using Garland et al.’s (1993) simulation-based approach, for example using the *aov*.*phylo* function implemented in “geiger” (Harmon et al. 2008). This approach uses the conventional (non-phylogenetic) MANOVA to fit the regression model and compute associated multivariate test statistics (e.g., the Pillai’s trace, Wilks’, Lawley-Hotelling, and Roy’s largest root tests; see a detailed review in Rencher 2002); the significance of the test is then assessed using simulations under the multivariate Brownian process (BM). To the best of our knowledge, maximum likelihood estimation of the multivariate PGLS regression model for testing the effect of continuous explanatory variables, pMANOVA for testing the effect of categorical independent variables (grouping of species), and pMANCOVA for testing the effect of categorical independent variables while accounting for the effects of other continuous variables (covariates), are not implemented in the popular R language. In high-dimensional settings (as the number of traits *p* approaches or exceeds the number of species *n*) the situation becomes even worst, as the evolutionary (or trait) variance-covariance matrix is singular, which prevents the use of likelihood-based techniques (Clavel et al. 2019).

Evolutionary biologists circumvent these current limitations for treating high-dimensional phenotypic datasets in two ways. A first approach consists in “reducing” the data to its principal component axes (PCs), after which the simulation-based approach on a reduced set of axes (p<n) and/or univariate phylogenetic regressions on individual axes are performed. The PC axes are obtained using either conventional (i.e. non-phylogenetic) or phylogenetic principal component analyses (PCA or pPCA, Revell 2009; Uyeda et al. 2015). One major limitation of these dimension reduction techniques is that using a restricted set of PC axes, or pPC axes from a misspecified evolutionary model, can lead to erroneous inferences (Uyeda et al. 2015). Until recently (but see Clavel et al. 2019 for recent developments that overcome these limitations), pPCAs in high dimension could only be computed assuming a Brownian model of trait evolution. Using such pPCs (or simply PCs) will likely be problematic in regression analyses when there are deviations from the underlying assumptions, although this has to our knowledge not been clearly assessed.

A second approach (Adams 2014; Goolsby 2016; Adams and Collyer 2018a; Collyer and Adams 2018) is inspired from distance-based pseudo-statistics initially developed in ecology to deal with cases when traditional parametric regressions cannot be performed (such as the PERMANOVA procedure used for count and abundance data, Anderson 2001; McArdle and Anderson 2001). In the phylogenetic distance-based approach (Adams, 2014, Goolsby 2016, Adams and Collyer 2018a; Collyer and Adams 2018), the trait data (both the response and the predictors) are first transformed using the inverse square-root of the phylogenetic variance-covariance matrix – assuming Brownian motion evolution on the tree; next, the sum of squared Euclidean distances between transformed data and predicted values under the regression model (and a simpler model representing the null hypothesis) are computed. Finally, a pseudo *F*-statistic is computed from the ratio of these sum of squared distances and compared to *F*-statistics computed on permuted data to assess statistical significance. While this approach can be performed on high dimensional phenotypic data, it in fact ignores trait correlations, as using Euclidian distances implicitly assumes that traits evolve independently (McArdle and Anderson 2001; Goolsby 2016). Indeed, the Euclidean distance approach sums the Sum of Squares (SS) across traits without accounting for Cross Products (CP) between traits. As a result, distance-based approaches using Euclidean distances are expected to be sensitive to unequal variances across the response variables and to the relationship between dispersion and effect change (e.g. the direction of variation across groups in relation to the main axis of variance) in the multidimensional space (Anderson 2001; McArdle and Anderson 2001; Warton et al. 2012). Using other distances – such as generalized or Mahalanobis distances (Mahalanobis 1936) – would avoid these issues, but would require inverting the evolutionary (traits) variance-covariance matrix, which would be possible only in low dimensional settings. In addition to these limitations, current implementations of the approach assume Brownian evolution and might be sensitive to departures from this assumption and/or to measurement errors that are likely to arise in large datasets.

Recently, we developed a framework, inspired from regularization techniques often used in conventional multivariate statistics (e.g., Friedman 1989; Hoffbeck and Landgrebe 1996; Warton 2008; Witten and Tibshirani 2009), for fitting a variety of trait evolutionary models (BM, Pagel’s lambda, Orstein-Uhlenbeck, Early Burst) to high dimensional phenotypic data (Clavel et al. 2019). Our approach uses penalized likelihood (e.g., Fan and Li 2001; Warton 2008; van Wieringen and Peeters 2016) to estimate evolutionary variance-covariance matrices under these models. Here, we build on this promising framework to develop phylogenetic regression techniques and associated tests (multivariate pGLS, pMANOVA, pMANCOVA) that can be applied to high-dimensional phenotypic datasets, while accounting for the amount of phylogenetic covariations actually present in the model residuals (e.g. by jointly estimating Pagel’s λ(Pagel 1999)). We implement this approach in the mvMORPH package (Clavel et al. 2015). We also implement similar functions to perform multivariate pGLS, pMANOVA and pMANCOVA models using maximum likelihood inference when *p<n*. We use extensive simulations to compare the performance of our approach to current alternatives, using the MANOVA as a case study. We illustrate the utility of the proposed method by analyzing the relationship between diet preferences and a geometric morphometric dataset describing mandible morphology in phyllostomid bats (Monteiro and Nogueira 2011). Previous analyses on a subset of PC axes suggested that phyllostomid bats have evolved into well differentiated ecomorphs (Monteiro and Nogueira 2011). Here we revisit these results using both our phylogenetic regularized MANOVA and the distance-based approach on the full multivariate superimposed Procrustes coordinates. Finally, we discuss some of the recent concerns in multivariate phylogenetic comparative methods and prospects for future work.

## MATERIAL AND METHODS

### Multivariate Phylogenetic Regressions and Associated Tests

We aim to test the potential effect of predictors (continuous and/or categorical) on *p* continuous traits from a comparative dataset comprised of a (ultrametric or non-ultrametric) phylogeny of *n* species (extinct or extant) with associated measured traits (mean trait value for each species). These tests can be performed by fitting generalized least squares (GLS) linear models of the form:

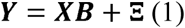

Where **Y** is the *n* ×*p* matrix of observed traits (response variables), **X** is a *n* ×*q* design matrix constructed from the observed predictors (e.g. each continuous predictor is encoded in one column in **X**, each categorical predictor is encoded in one or several columns using dummy variables, and there can be an additional column of 1 for the intercept), **B** is a *q* × *p* matrix of unknown coefficients describing the dependencies of traits to the predictor variables (slopes and intercepts for continuous predictors and mean traits for categorical predictors),and Ξ is a *n*×*p* matrix of errors distributed according to a matrix-normal distribution ℳ 𝒩_*n,p*_(0,***C***,***R***) (Gupta and Nagar 1999). **C** is the phylogenetic variance-covariance matrix which represents expected variances and covariances and depend on the phylogeny and an evolution model (Rohlf 2001; Felsenstein 2004), and ***R*** is the *p × p* thevariance-covariance matrix of residuals under the regression model.

Once estimates 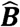 Ĉ and 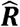 of such linear models are obtained (see below), common multivariate tests (e.g., the Wilks’s Λ, Pillai’s trace, Lawley-Hotelling, and Roy’s largest root tests Rencher 2002) are constructed from statistics based on the eigenvalues *d*_1_,…,*d*_*s*_ of the *p ×p* matrix **E**^**–^1^**^***H*** where ***E*** is the error (or residual) matrix, ***H*** is the hypothesis matrix (which measures the deviation of predicted response values under the complete and null hypothesis models), and *s* is the rank of **E**^**–1**^***H*** (Rencher 2002; Huberty and Olejnik 2006; Fox 2015). For instance, Wilk’s Lambda test statistic is defined as (Rencher 2002; Fox 2015):

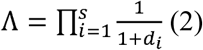

***E*** (also called the residual Sums of Squares and Cross Products (SSCP) matrix) is given by 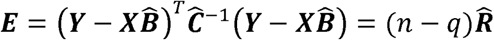 and ***H*** (also called the hypothesis SSCP matrix) is given by 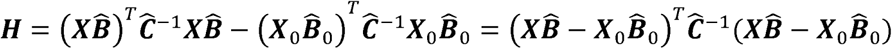 Where ***X*** _0_ is the design matrix corresponding to the null hypothesis and 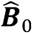 is the corresponding matrix of parameter estimates. Other ***H*** matrices corresponding to different null hypotheses (as in the case of general linear hypothesis testing) can be formulated and treated in the same way (Supplementary Material).

### Maximum Likelihood and Penalized Likelihood Regressions

Estimates of ***B, C***, and ***R*** can be obtained by maximizing the following (restricted) log-likelihood (see Clavel et al. 2019 and Appendix 1):

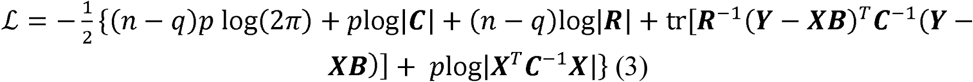

Where for a given matrix **A** tr (**A**) stands for the trace (the sum of the diagonal elements), |**A**| for the determinant, ***A***-1 for the inverse, and **A**^T^ for the transpose.

The maximum likelihood estimate of ***B*** is given by (Rao and Toutenburg 1999; Timm 2002):

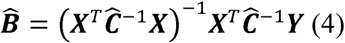

And the (restricted) maximum likelihood estimate of ***R*** is given by (Searle et al. 1992; Rao and Toutenburg 1999):

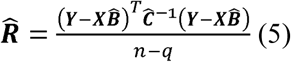

### Maximum likelihood

When *p* ≤*n–q*, the inverse and the log determinant of 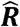 can be computed and we find Ĉ, 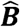, and 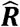 by maximizing the likelihood (Equation 3) (e.g. Hansen and Martins 1996; Pagel 1999; Blomberg et al. 2003; Felsenstein 2004; Clavel et al. 2015, 2019; Manceau et al. 2017). In this case, it is straightforward to invert ***E***, and thus to compute ***E***^*–1*^***H*** and various test statistics. We assess the significance of the test following the procedure used in conventional multivariate linear regressions (OLS): we transform the various test statistics to an approximate *F* statistic, which significance is assessed by comparison to the *F*-distribution with appropriate number of degrees of freedom (Rencher 2002, chapter 6).

### Penalized Likelihood

When the number of traits *p* approaches (or equals) *n*-*q*, 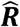 does not necessarily provide a good estimation of ***R***; when *p* exceeds *n-q*, 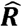 is singular and we can no longer compute its inverse nor the logarithm of its determinant (James and Stein 1961; Witten and Tibshirani 2009). In this case, we cannot directly compute **E**^−1^ and the test statistics. We circumvent these issues by building upon our recently developed penalized-likelihood approach (Clavel et al. 2019). In Clavel et al. (2019), we provide several regularized estimators of ***R***. Here we use the popular linear shrinkage estimator, which is given by (Equation 3 in Clavel et al. 2019):

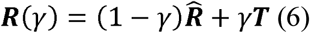

where γϵ[0,1] is a regularization (or tuning) parameter that controls the amount of shrinkage from 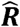 to a *p* by *p* target matrix ***T***. We chose this estimator as it is the fastest to recent studies showed that it provides good performances with conventional multivariate statistics (e.g., Tsai and Chen 2009; Engel et al. 2015; Ullah and Jones 2015). Here we take the matrix ***T*** to be diagonal with all diagonal elements equal to 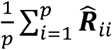. Hence, the limit γ=1 corresponds to the case when the traits are assumed to be independent and to share the average trait variance. We chose this target matrix as it is rotation invariant (proportional to the identity matrix, Clavel et al. 2019); preliminary simulation analyzes showed that this shrinkage provided a good compromise between precision in the estimation of ***R*** and computation time (results not shown). Following Clavel et al. (2019), we find the optimal value for γ and the estimator ***R*** (γ) by maximizing the leave-one-out cross-validated (LOOCV) log-likelihood corresponding to a “ridge” penalized log-likelihood (Appendix 1). Inverting ***R*** (γ) provides us with a regularized estimate of from which the test statistics are computed. Finally, we approximate the distribution of a given test statistic using a permutation procedure, following the general approach of permuting the residuals under the reduced model (Freedman and Lane 1983; Anderson and Braak 2003) and borrowing ideas bootstrapping procedures (e.g., Pennell et al. 2015; Khabbazian et al. 2016). We simulate several (999 in what follows) new datasets 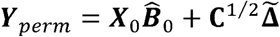 where each 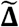 is from a matrix of permuted residuals obtained by permuting the rows of 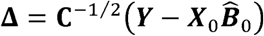, and compute the test statistic corresponding to each of these datasets using our penalized likelihood procedure. This procedure provides an approximate distribution of the test statistic which is used to assess statistical significance (see also Hall and Wilson 1991 for general guidelines on non-parametric estimation of null distribution). In addition to this procedure, where γ, ***C, B***, and ***R*** are optimized by maximizing the LOOCV on each of the permuted datasets, We consider a more efficient (approximated) procedure that consists in first transforming the “tests” and “training” samples used in the LOOCV, similar to the one used by Warton (2008). This approach (detailed in Appendix 2) is approximate as it assumes that ***C*** to be the penalized likelihood estimate obtained from the empirical data.

### Implementation

We implemented the fit of the multivariate phylogenetic least squares in the *mvgls* function in the R-package mvMORPH publicly available on CRAN (Clavel et al. 2015) and on gitHub (https://github.com/JClavel/mvMORPH). The option “method” allows a user to fit the linear model either by likelihood (when *p* ≤ *n* – *q*, “LL”) or by penalized likelihood (“LOOCV”). The linear hypothesis tests are implemented in a separate function called *manova.gls*. This function takes as input an object of class “mvgls”, output of the *mvgls* function, and allows performing the parametric (when *p* ≤ *n* – *q*,) as well as the (exact and approximate) permutation-based hypothesis testing. The user can also perform general linear hypothesis testing by specifying specific matrices in the “L” (contrasts coding matrix) and “rhs” (right-hand side matrix) options (Supplementary Material). We also offer the possibility to perform parallel calculus (forking) using the base package “parallel” in R to compute the permutation-based distribution of the test statistic. The functions *mvgls* and *manova.gls* are structured as the *lm* and *manova* functions from the *stats* base R-package (R Development Core Team 2016).

In our simulations, we used Pagel’s λ phylogenetic model for computing **C** (Pagel 1999), with a unique lambda common to all traits. This phylogenetic model is useful from a statistical point of view as it can be seen as a mixed model where both a Brownian motion diffusing along the branches of the tree and random independent noise contribute to interspecies trait variation (Housworth et al. 2004, see also the Supplementary Material in Clavel et al. 2019). This flexibility should allow accommodating a range of phylogenetic signal compared to methods that rely exclusively on Brownian motion and has been shown to reduce the risk of model misspecification in univariate phylogenetic regressions (Revell 2010). As test statistics, we implemented Wilks’s λ, Pillai’s trace, Hotelling-Lawley and Roy largest root test (Hotelling 1931; Wilks 1932; Lawley 1939; Roy 1953; Pillai 1955). Pros and cons of these statistics depend on the multidimensional structure of the data and are discussed at length in the literature (e.g., Olson 1974; Rencher 2002). In what follows, we considered the Wilks Λ statistic with a significance level, *α*= 0.05. We chose this statistic as it is equivalent to computing a likelihood ratio (Wilks 1932). Finally, for multiple regressions and factorial designs, we implemented three different ways to test for the effect of a specific variable while accounting for the effect of others. If the “type” option in the *manova.gls* function is set to I, a sequential decomposition of the SSCP is performed; if it is set to II or III, a marginal or partial decomposition is performed. Which type of decomposition to use depends on the question at stake and has been discussed elsewhere (Langsrud 2003; Heiberger and Holland 2015; McFarquhar 2016).

### Testing the performance of the PL GLS: the MANOVA as a case study

We used simulations based on the one-way MANOVA procedure for a binary predictor variable (Rencher 2002; Huberty and Olejnik 2006) to compare the performances of the proposed multivariate approach with current alternatives. We considered the following approaches (Table 1): i) the simulation-based MANOVA (Garland et al. 1993) implemented in the *aov.phylo* function in geiger (Harmon et al. 2008), ii) the likelihood-based GLS-MANOVA we implemented here in mvMORPH (function *manova.gls*, option “LL”), iii) the distance-based MANOVA (Adams 2014) implemented in the *procD.pgls* function in geomorph version 3.0.7 (Adams and Otarola-Castillo 2013; this function is equivalent to the *lm.rrpp* function from the RRPP package, Collyer and Adams 2018); note that earlier versions of *procD.pgls* had an error in the permutation procedure that lead to very high type I errors (results not shown) and iv) the regularized PL-MANOVA we developed here (with both the exact and the approximated permutation strategies). We also used the simulation-based MANOVA on datasets that were reduced to their three first PC (*prcomp* in R) and phylogenetic pPC axes (*phylo.pca* in phytools, Revell 2009; Uyeda et al. 2015). Finally, we performed as a benchmark the conventional (non-phylogenetic) MANOVA based on ordinary least-squares (Rencher 2002) using the *lm* and *manova* R base functions.

**Table 1.** Summary of available phylogenetic approaches for performing regressions with multivariate trait data and associated propertie

| Approach | Predictor | Limited to BM | Limited to $p < n$ | Assumes uncorrelated traits |
| --- | --- | --- | --- | --- |
| distance-based<sup>1</sup> | All | Yes | No | Yes |
| simulation-based<sup>2</sup> | Only categorical | Yes | Yes | No |
| regressions on (p)PCs | All | Yes | No | No |
| ML-MANOVA<sup>3</sup> | All | No | Yes | No |
| PL-MANOVA<sup>3</sup> | All | No | No | No |

<sup>1</sup> the distance-based approach is implemented in geomorph (function *procD.pgls*) and in the "RRPP" package (function *lm.rrpp*)
<sup>2</sup> the simulation-based approach is implemented in geiger (function *aov.phylo*).
<sup>3</sup> new approaches implemented in mvMORPH (function *mvgl*s and *manova.gls*)

In our simulations we fixed the number of species to *n* = 32 and varied the number of traits *p* from 5 to 100 (*p*=5, 15, 25, 31, 50, 100). We simulated phylogenetic trees under a pure-birth process, scaled to unit height, using the *pbtree* function in “phytools” (Revell 2012). We considered three different values of *λ*, held identical across traits (*λ* = 0,0.5,1). These values represent scenarios ranging from traits showing no phylogenetic signal (*λ*= 0) to traits having evolved by Brownian motion (*λ*= 1). We simulated random covariance matrices ***R*** with varying degree of average correlation between traits (*ρ* = 0.2,0.5,0.8): we sampled matrices from a Wishart distribution with degree of freedom fixed to *df* = 100 + *n* and parameter matrix Σ= ***R***_*ρ*_ /*df*, where ***R***_*ρ*_ is a *p* by *p* matrix with off-diagonal entries *ρ* and diagonal entries 1 using the *rwishart* function in the “dlm” package (Petris 2010). In comparison with previous approaches (e.g. in Uyeda et al. (2015) and Clavel et al. (2019)), this approach allows controlling for the average amount of correlation between traits. We simulated trait data of size *n* by *p* on the transformed trees (according to Pagel’s *λ* values) under a multivariate Brownian motion with covariance matrix ***R*** using the *mvSIM* function in “mvMORPH” (Clavel et al. 2015).

In order to assess the type I and type II errors of the MANOVA test, we considered two balanced species groups of size *n* = 16 each, with either differences in mean trait values (***μ***_1_=0, ***μ***_2_ ≠ ***μ***_1_, Fig.1a,b,c) or no difference in mean trait values (***μ***_1_ =0 and ***μ***_2_ = ***μ***_1_, Fig.1d). Following previous studies (Warton 2008; Ullah and Jones 2015) we considered three alternative scenarios to the null hypothesis (*H*_0_: ***μ***_1_= ***μ***_2_ : we shifted the mean for the second group such that the differences between groups are expressed i) along the main axis of variance (Fig 1a, shift along the first eigenvector of ***R*** 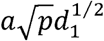 units), ii) along the second axis of variance (Fig 1b, shift along the second eigenvector by 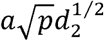 units), and iii) along all axes (Fig 1c, shift along each eigenvectors by 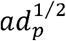 units), where *d*_1_, *d*_2_,..,*d*_*p*_ are the eigenvalues of ***R*** and *a* is fixed at 0.3 (0.6 for convenience in the first scenario; see details of the procedure in the Supplementary Material).

**Figure 1.**
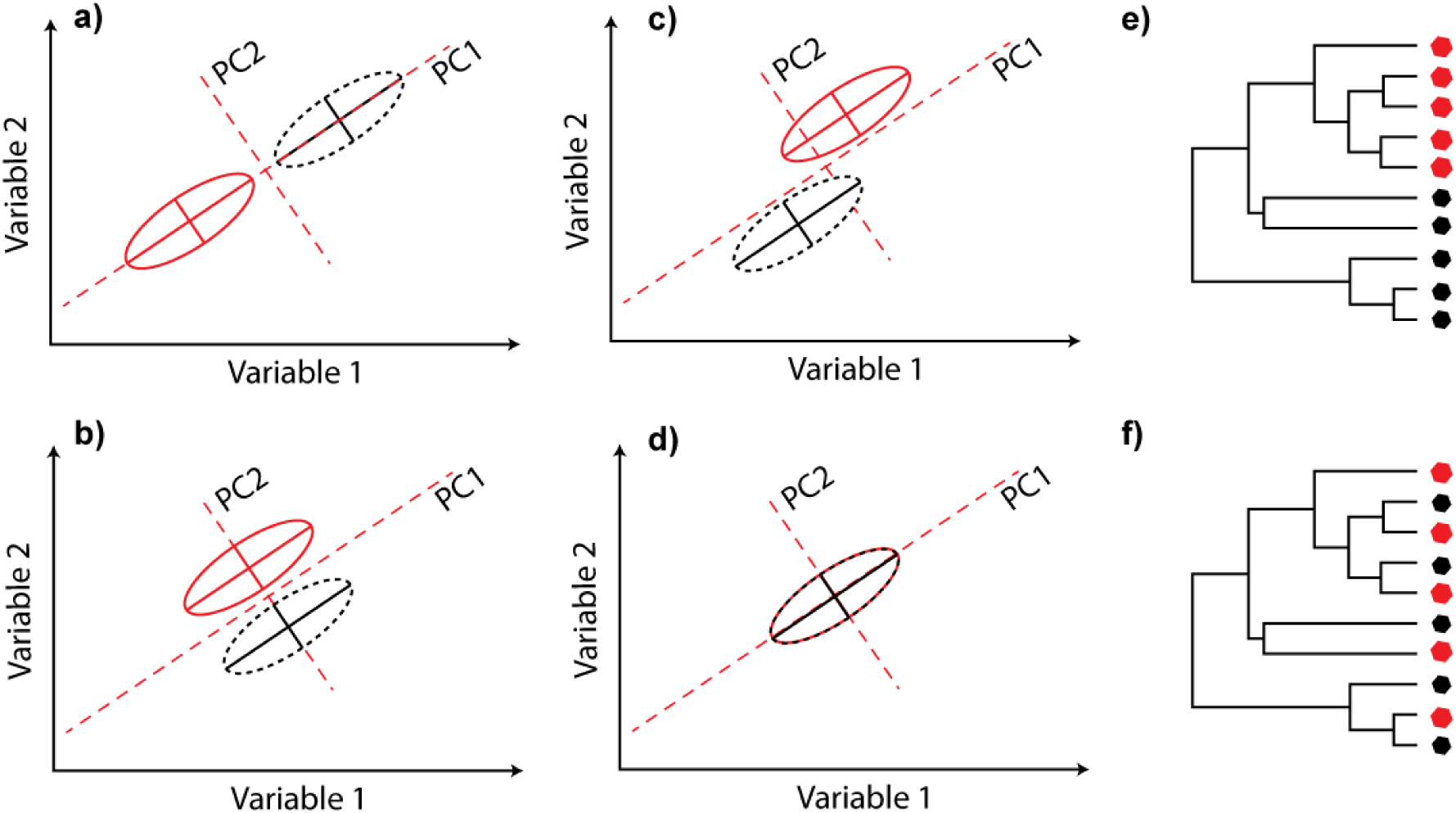
Schematic illustration (with a bivariate trait) of the various scenarios considered in our simulations. The black and red colors represent the two species groups. (a, b & c) Main trait differences between the two species groups occur along (a) the main axis of variance, (b) the second axis of variance, or (c) both axes. Differences between groups such as those described in (b) and (c) require truly multivariate tests, as differences are difficult to detect on each variable taken separately (i.e., on the marginal analyses). In (d), there is no difference between the two species groups (i.e., the null hypothesis of the MANOVA test). (e & f) the distribution of the predictor binary state (which determines the species groups) is (e) phylogenetically clustered or (f) uniformly distributed.

We also considered two alternative scenarios for the distribution of the binary predictor with respect to the phylogenetic relationships (Fig 1e,f). In the first scenario, the first 16 species (in the *tip.label* “ape” phylo object in R) were assigned mean vector ***μ***_1_, and the 16 last species the mean vector ***μ***_2_. Given that pure-birth trees tend to produce balanced topologies, this first strategy imposes a strong phylogenetic structure in the distribution of the binary predictor (Fig 1e). In the second scenario, the group labels were alternated across consecutive tips leading to a more dispersed distribution of the binary state predictor on the tree (Fig 1f).

We simulated 1000 datasets for each parameter set [*p*=(5, 15, 25, 31, 50, 100), *λ*= (0,0.5,1) and *ρ* = (0.2,0.5,0.8)] and scenario [no difference, shift along the first, second, or all eigenvectors, and phylogenetic structure of the binary predictor]. Finally, we performed the various tests on each dataset and recorded the number of datasets for which a significant difference between groups was found. When *p* ≤ *n* – *q*, this resulted in 8 tests per dataset. When *p* ≥ *n* – *q*, only the distance-based MANOVA, the simulation-based MANOVA on PC (and pPC) axes, and the regularized PL-MANOVA were performed (4 tests). Importantly, our GLS-MANOVA (when *p* ≤ *n* – *q*) and PL-MANOVA (when *p* ≤ *n* – *q*, or *p* ≥ *n* – *q*,) estimate phylogenetic signal in the model residuals using Pagel’s *λ* when performing the tests. The other tests assume either no phylogenetic signal (conventional MANOVA based on OLS), or Brownian motion (the distance-based and simulation-based approaches, as well as approaches using pPC axes computed using *phylo.pca*). All the analyses were performed on a Linux platform with R version 3.4.4 (R Development Core Team 2016).

### Empirical Illustration: the evolution of the mandible in phyllostomid bats

We applied our PL-MANOVA on a high-dimensional comparative dataset (*p* ≥ *n* – *q*) describing mandible form (size and shape) for 49 species of phyllostomid bats (leaf-nosed bats) with respect to their diet. We used the phylogenetic tree and mandible data from Monteiro and Nogueira (2011) which consists of 11 anatomical landmarks and 25***Figure 3.*** *Comparison of the statistical performances (statistical* semilandmarks for 2D coordinates (aligned by Procrustes Generalized Analysis; see Figure 2), as well as the logarithm of the condylobasal length used as a proxy for size (totaling 73 variables forming a size-shape space; Mitteroecker et al. 2004). In their original study, Monteiro and Nogueira (2011) used five different diet categories (frugivores, insectivores, carnivores, nectarivores, and sanguivores). We applied our PL-MANOVA (function *mvgls* and *manova.gls* in “mvMORPH” using Pagel’s *λ*) to test whether mandible form depends on diet, using four different species groupings (Figure 2): i) two species groups corresponding to feeding modes that involve a considerable amount of mastication (insectivores, carnivores and frugivores) or little or no mastication (sanguivores, nectarivores); ii) three groups obtained from the previous ones but treating sanguivores and nectarivores separately; iii) four groups further separating frugivores from animalivores (i.e. insectivores and carnivores) iv) the five original groups. We compared the results to those obtained with the distance-based MANOVA approach (function *procD.pgls* in “geomorph”). We further illustrate the application of general linear hypothesis testing by assessing, in the five groups scenario, whether there was a significant difference between carnivores and insectivores, between sanguivores and nectarivores, and between frugivores and animalivores.

**Figure 2.**
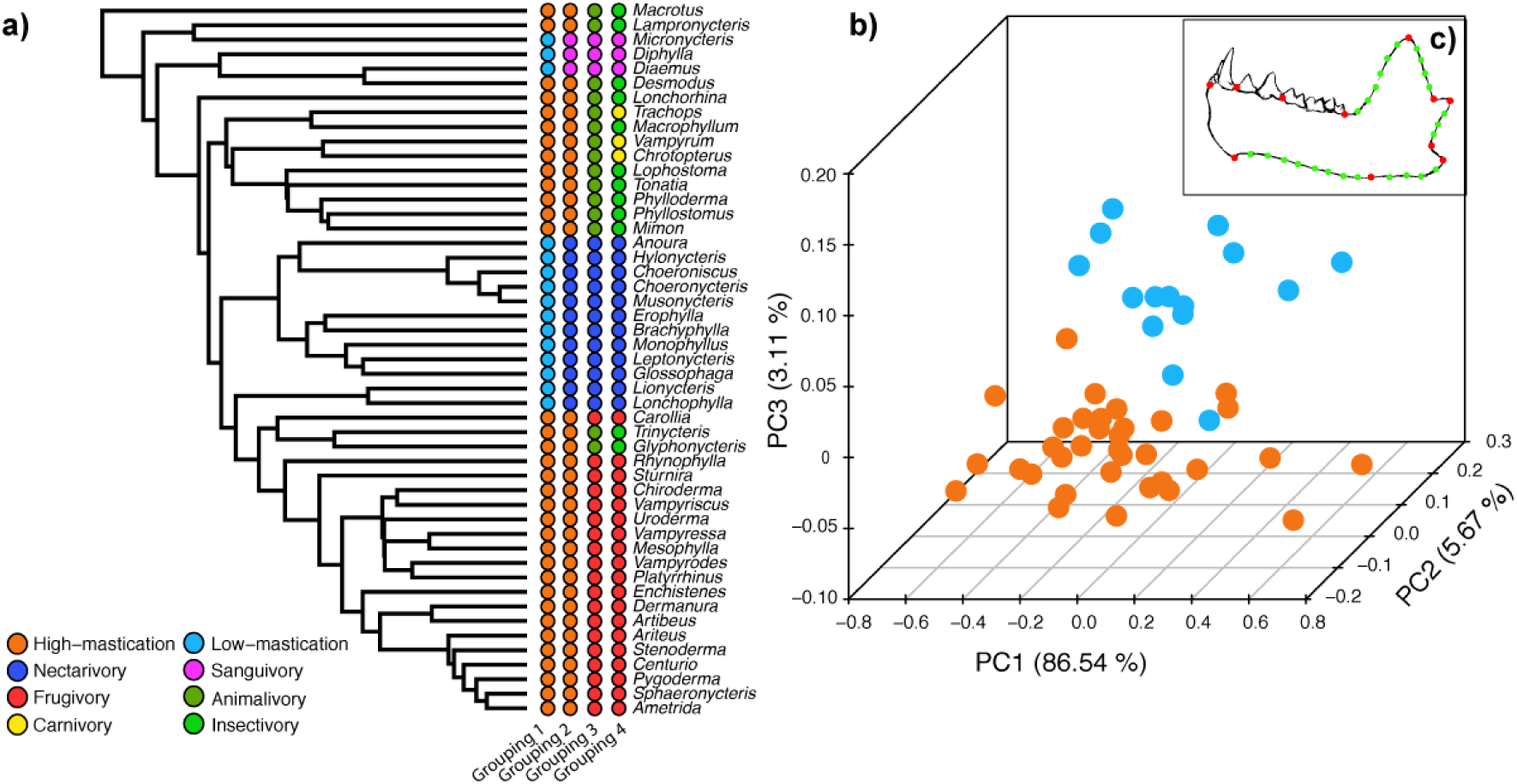
The phyllostomid bats dataset. a) the phylogenetic tree of phyllostomid bats with four different species grouping reflecting diet preferences and feeding modes. b) Scatter plot of the three first phylogenetic PC axes obtained using the mvgls.pca function in “mvMORPH” (PL estimate of λ = 0.45). Species are colored according to Grouping 1 (high mastication in orange, low-mastication in blue). The two groups are well separated on the third axis of variation of the size-shape dataset. c) Illustration of the landmarks (in red) and semilandmarks (in green) used for describing the mandible shape in Monteiro and Nogueira (2011).

**Figure 3.**
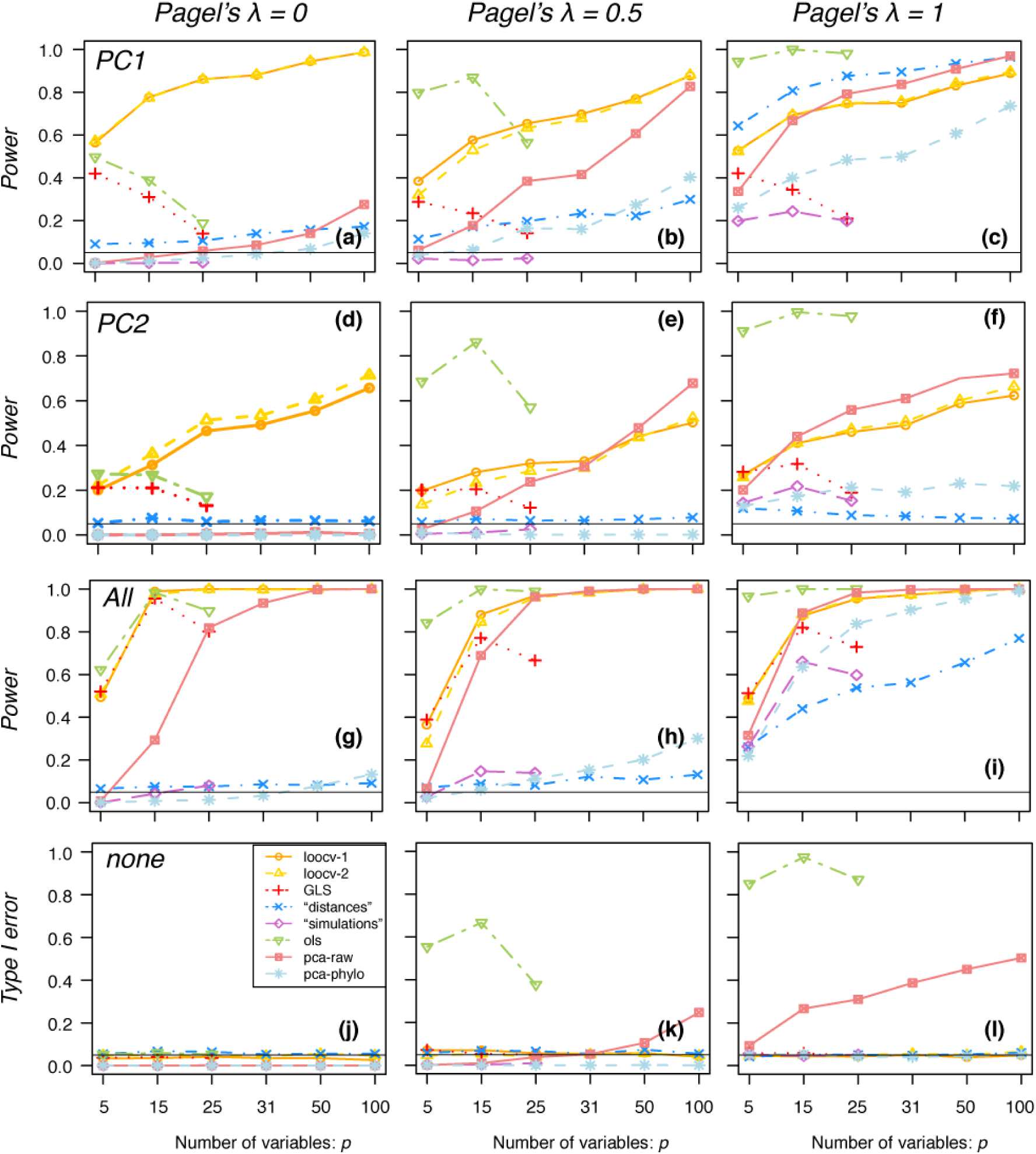
Comparison of the statistical performances (statistical power and Type I error) for the various MANOVA approaches with a phylogenetically clustered binary predictor variable. The three columns correspond to simulations with no phylogenetic signal (left column, lambda=0), intermediate phylogenetic signal (middle column, lambda=0.5), and BM (right column, lambda=1). On the top row (a, b, c) the differences between groups occurred along the first axis of variance (PC1); on the second row (d, e, f), they occurred along the second axis (PC2); on the third row (g, h, i) they occurred along all axes (All). On the last row (j, k, l) there were no differences between groups (none). “loocv 1” refers to the PL-MANOVA with exact permutation procedure and “loocv-2” to the approximated approach; “GLS” refers to the maximum-likelihood MANOVA based on generalized least squares; “distances” refers to the distance-based approach implemented in geomorph; “simulations” refers to the simulation-based approach implemented in geiger; “OLS” refers to the conventional (non-phylogenetic) ordinary least squares MANOVA; “pca-raw” and “pca-phylo” refer to the simulation-based MANOVA on the three firsts PC axes and phylogenetic PC axes (obtained from phylo.pca in phytools).

## RESULTS

All the results shown in the main text correspond to datasets simulated with an average correlation among traits (*ρ*= 0.5). Results with low (*ρ*= 0.2) and high (*ρ*= 0.8)average correlations are shown in the Supplementary Material.

### Statistical Power and Type I Error

Our PL-MANOVA tests (“loocv-1” and “loocv-2” approaches in Figs. 3-4 and S1-4) perform well under all types of differences between groups (differences along the first, second, or all axes of variation), and for the various amount of phylogenetic signal (Pagel’s λ), regardless of whether the binary predictor is phylogenetically clustered (Fig. 3 and S1, S3) or not (Fig. 4 and S2, S4). Their power to detect differences between groups increases with the number of traits, and their type I error rate is always at their nominal level (0.05). The efficient approximated test (“loocv-2”) performs as well as the more computationally demanding exact test (“loocv-1”).

**Figure 4.**
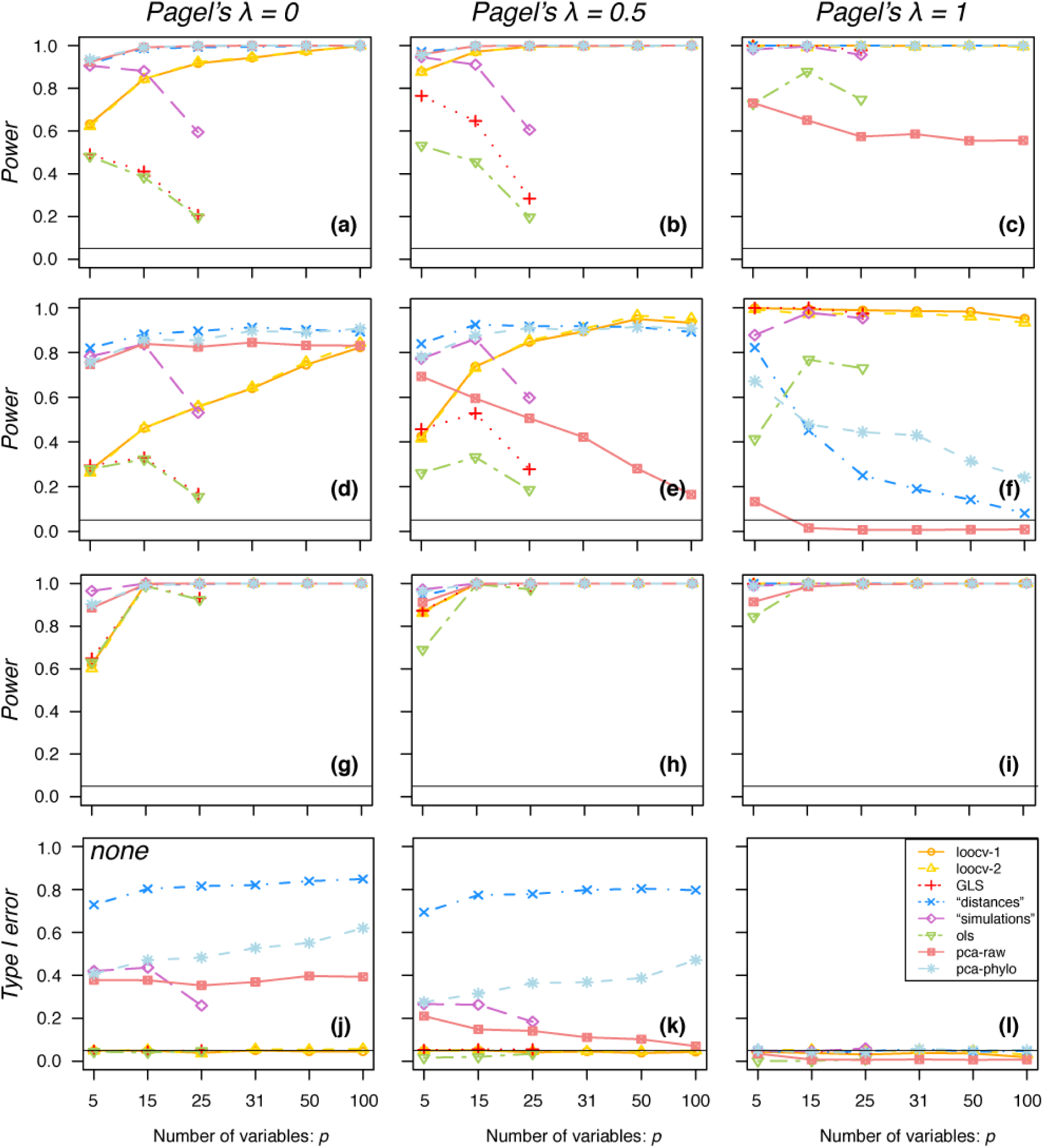
Comparison of the statistical performances (statistical power and Type I error) for the various MANOVA approaches with a non-phylogenetically clustered binary predictor. Notations are as in Figure 3.

The performances of the other tests depend on the data. The GLS-MANOVA test performs as well as the PL-MANOVA when the number of traits is low, but rapidly loses power as the number of traits increases. As expected, the OLS performs as the GLS-MANOVA in the absence of phylogenetic signal (*λ* = 0); however it has a high type I error in the presence of phylogenetic signal (*λ* > 0) when the binary predictor is phylogenetically clustered (Fig 3 k,l). The simulation-based approach generally has a lower power than the GLS-MANOVA and PL-MANOVA approaches, and particularly so when there are deviations from the Brownian process (*λ* < 1) and when the binary predictor is phylogenetically clustered (Fig. 3 a,b,d,g,h); in addition, when *λ* < 1 and the binary predictor is not phylogenetically clustered, it is subject to a high type I error (Fig. 4 j,k). Similarly, the distance-based approach has a low power when *λ* < 1 and the binary predictor is phylogenetically clustered (Fig. 3 a,b,d,e,g,h), and it is subject to a high type I error when *λ* < 1 and the binary predictor is not phylogenetically clustered (Fig. 4 j,k); in addition, it has a low power even under the Brownian process (*λ* = 1) when the differences between species groups are not located on the first axis of variation (Fig. 3f,i, and 4f; Fig. S3-4). Finally, dimension reduction techniques (“pca-raw” and “pca-phylo”) have high type I errors if there is an unaccounted-for phylogenetic signal and the binary predictor is phylogenetically clustered (“pca-raw” when, *λ* > 0, Fig. 3k,l), or if there are deviations from the Brownian process and the binary predictor is not phylogenetically structured (“pca-raw” and “pca-phylo” when, *λ* < 1, Fig. 4j,k).

Overall, the performances of the different approaches do not strongly depend on the degree of among-traits correlations (Figs 3-4 and Figs. S1-S4). The main exception is the distance-based approach, which has a decreasing power to detect differences between groups with increased correlations among traits when differences are not located on the first PC axis (Fig. S3-4, second row). When differences between groups occur on the first axis of variation, most approaches show higher power when the correlation among traits is low (compare the top row in Fig. S1-2 vs Fig. S3-4). In contrast, when the simulated differences are spread across all the PC axes, the power is higher with high among traits correlations (compare the third row in Figs. S1-2 vs S3-4).

### The evolution of the mandible in phyllostomid bats

Our PL-MANOVA tests revealed a significant difference (i.e. rejected *H*_*0*_) on mandible morphology in phyllostomid bats, for the four diet grouping schemes considered (Table 2). The estimated phylogenetic signals in the residuals of the models were rather low (Pagel’s *λ* = 0.44 − 0.08; Table 2a) and estimated average absolute correlations between variables ranged from 0.32 to 0.39. In comparison, the distance-based approach did not detect a significant difference in any grouping scheme (Table 2b). To assess whether this discrepancy was related to the Brownian motion assumption of the distance-based approach while there was a low phylogenetic signal estimated in the residuals of the models, we reconducted the analysis after rescaling the phylogenetic tree according to the PL estimates of Pagel’s λ. In this case the scheme with five diet groups appeared significant, but not the others (Table 2b). To evaluate whether the discrepancy for the three first groupings was related to differences across groups occurring on the second and/or higher axes of variation rather than the first axis, we represented the differences among groups visually by computing pPC axes (Figs. S5-6, see also Fig. 2b) and performed univariate ANOVAs on the three firsts pPC axes (see Supplementary Material). Indeed, differences among groups were visible in the three first groupings, but they occurred along the second and/or third axes, not the first one (Fig. S5-6 and Table S1). In the analysis on the five diet groups, on the other end, some slight differences occurred on the main axis between carnivorous and insectivorous bats (although the number of carnivorous species is low). The main axis mostly displays changes in sizes, with carnivorous bats that tend to be slightly larger than insectivorous ones (Monteiro and Nogueira 2011). Our general linear hypothesis tests confirmed a significant effect of carnivory and insectivory on mandible morphology. They also detected a significant effect of nectivory versus sanguivory on mandible morphology, even though these two diets have low masticatory demands, as well as between frugivory and animalivory, even though these two diets involve a substantial amount of mastication (Table S2).

**Table 2.** Effects of diet on the mandible in phyllostomid bats. **(a)** phylogenetic signal estimated by the PL-MANOVA and (b) results of the MANOVA tests.

| a) | Estimated phylogenetic signal (by PL) in the residuals of the model |  |  |  |
| --- | --- | --- | --- | --- |
|  | Grouping 1 | Grouping 2 | Grouping 3 | Grouping 4 |
| Pagel's $\lambda$ | 0.44 | 0.42 | 0.32 | 0.08 |

| b) | Tests significance (p-values) |  |  |  |
| --- | --- | --- | --- | --- |
| PL-MANOVA | **0.001** | **0.001** | **0.001** | **0.001** |
| distance-based MANOVA | 0.509 | 0.585 | 0.689 | 0.056 |
| distance-based MANOVA* | 0.236 | 0.206 | 0.066 | **0.001** |

\*phylogenetic tree transformed according to the PL estimates of $\lambda$ in (a)

## DISCUSSION

We developed new regularized multivariate statistics and associated tests (PGLS, phylogenetic MANOVA and MANCOVA) for assessing the effect of continuous and/or categorical predictors on high-dimensional phenotypic datasets in a phylogenetic context. We focused on the MANOVA to test the performance of these tests using simulations and expect the performance of the other tests to be similar, as they rely on the same penalized-likelihood framework. The penalized-likelihood tests outperform current alternatives in both low (*p*≤ *n*-*q*) and high (*p*≥ *n*-*q*) dimensions. They are more powerful for detecting differences among groups, in particular when these differences do not only occur on the first axis of variation. By estimating the amount of phylogenetic signal from the data, they also avoid false positives that can occur when ignoring phylogenetic signal (e.g. using non-phylogenetic multivariate statistics) or when overestimating this signal (e.g. assuming a Brownian structure in the residuals). Applying these new methods to phyllostomid bats, we found a significant effect of diet on mandible morphology that would not have been detected by other methods.

We highlighted two main aspects of multivariate comparative datasets that impact the ability of current statistical tests to detect the effect of predictors on trait data and the Type I error rate associated with these tests. First, the direction of the variation matters (Fig. 1); previous studies that have investigated the statistical performances of phylogenetic multivariate linear regressions or MANOVA (Goolsby 2016; Adams and Collyer 2018a, b) have focused on scenarios where differences between groups or relationships between explained and explanatory variables are expressed along the main axis of variation (e.g. Fig. 1a). However, as we have illustrated here with the phyllostomid bats, differences of biological interest may not occur along this first axis only. And as we have shown with simulations, comparative methods may perform differently depending on the axis along which variations occur. Hence, it is important to test multivariate comparative methods under various designs (e.g. Fig. 1b,c). When evaluating the performance of the distance-based approach (Adams and Otarola-Castillo 2013; Collyer and Adams 2018), we found that it had a very low power (up to the baseline nominal level of 5%) to detect differences between groups when these differences were not located on the main axis of variation, in particular when traits were moderately or highly correlated. These methods are based on Euclidean distances, which implicitly assume that the variables are uncorrelated with equal variances and are known to be sensitive to the mean-variance (or location-dispersion) effect (e.g., Anderson 2001; Warton et al. 2012). When traits are highly correlated, the first axis of variance explains a large part of the total variation, and differences between groups along the other axes are not detected by distance-based approaches. Penalized-likelihood approaches (as well as conventional multivariate approaches), on the other hand, are able to detect such differences. In the phyllostomid bats, for example, the PL approach detected differences that occurred on axes other than the first axis, while the distance-based approach did not.

Second, the phylogenetic structure of the data matters. Most available multivariate phylogenetic linear models are restricted to the Brownian motion process. However most empirical datasets will likely deviate from this BM assumption. In the phyllostomid bats dataset for example, we found a relatively low phylogenetic signal in the residuals of the different models, even though we could have expected a rather strong signal given that the group is thought to have experienced an adaptive radiation. More generally, high-dimensional datasets likely accumulate measurement errors and uncertainties (Goolsby et al. 2017) that may also lead to departures from the BM assumption. We have shown with simulations that methods relying exclusively on the BM process can have low power and/or high type I error rates when departures from the BM assumption occur. In the phyllostomid bats, differences associated with carnivory on the condylobasal length (a proxy for body-size) were not detected by the distance-based approach, even though these differences were observed along the main axis of variance, unless the tree was rescaled according to the corresponding PL estimate of phylogenetic signal. The PL approaches did detect these differences. Similarly, conventional (non-phylogenetic) methods have low power and/or high type I error rates when there is in fact a phylogenetic signal in the data. As we have no way to assess the phylogenetic signal – by which we mean the phylogenetic correlation structure – in the residuals *a priori* (i.e. without fitting a model), it is not possible in practice to know if and how much the data deviates from the BM or no-signal assumptions. By allowing an estimation of the signal in the residuals jointly with the model parameters, our approach accommodates the various data types with low type I error and good power, as was previously shown for univariate linear regressions (Revell 2010).

The phylogenetic structure of the predictors also influences the performance of the multivariate tests, although in a complex way. Adams and Collyer (2018b) argued that phylogenetic clustering of the predictors reduces the statistical power of multivariate tests, and suggested to perform prior tests of phylogenetic signal on the predictor variables in a systematic way – e.g. using two-block partial least squares 2B-PLS. However, we found that the effect of phylogenetic structure in the predictors on statistical performance depended on the phylogenetic structure in the residuals (see also Revell 2010). For example, we found high type I error rates with a non-clustered binary predictor when there were departures from the BM assumption in the residual structure. Hence, conditional tests based on measured phylogenetic signal in the predictors (or the responses) will be ineffective in practice and potentially misleading, as has been discussed before (Rohlf 2006; Revell 2010). In addition, we found that the power of GLS-MANOVA and PL-MANOVA when there was a strong phylogenetic signal in the predictor was not reduced compared to conventional methods used in their optimal conditions (i.e. when there is no signal in the residuals and *p*<*n*). Hence, multivariate tests can be performed with satisfactory power and type I error rates even when the predictors are phylogenetic clustered, provided phylogenetic structure in the residuals is properly accounted for; our regularized approaches allow doing so.

Evolutionary biologists often reduce high-dimensional datasets using Principal Component Analyses before performing multivariate tests (e.g. simulation-based MANOVA) using either conventional PC axes or phylogenetic PC axes assuming Brownian evolution. We have shown that this approach has a reduced power and is prone to high type I error rates, regardless of whether conventional or pPC axes are used. Conventional PC axes are sometimes favored as they are more straightforward to interpret biologically (Polly et al. 2013). This approach is known to affect model selection (Uyeda et al. 2015), and we have shown here that it also generates a lot of false positives in multivariate tests when there is a phylogenetic signal in the residuals. Phylogenetic PCA can account for such phylogenetic signal (Revell 2009), but it requires the phylogenetic model to be known prior to data reduction (Uyeda et al. 2015). We have shown that when the evolutionary model is misspecified (e.g., phylogenetic PCA based on Brownian motion is used when there is no or little phylogenetic signal), multivariate tests performed on pPCs axes are prone to high type I error rates. We therefore advise avoiding data reduction techniques that make strong *a priori* hypotheses on the phylogenetic structure (i.e. no structure or BM structure), and instead estimating the phylogenetic structure while performing the tests; the penalized likelihood tests proposed here allow doing this and are not prone to high type I error rates.

Several directions can be envisioned to further improve multivariate phylogenetic regression tests. For example, we estimated phylogenetic signal using Pagel’s λ model, which can be viewed as a phylogenetic mixed model (PMM) combining a Brownian motion process with independently and normally distributed errors (Housworth et al. 2004; Clavel et al. 2019). Even more flexible PMMs could be envisaged, for example based on the Ornstein-Uhlenbeck process (Mitov and Stadler 2018); they are already available in the penalized-likelihood framework (Clavel et al. 2019). Future improvements should also consider models where each trait may have its own level of phylogenetic signal – as was done in lower dimensional settings (e.g., Freckleton 2012; Ho and Ané 2014) – rather than assuming a common (or average) signal across trait as we did here. In addition, we considered a common covariance for each group, as is assumed by standard (M)ANOVA procedures which use a pooled estimate of the covariance matrix (Rencher 2002). Methods that account for group-specific covariances should probably be preferred when there are departures from this assumption. Such methods have already been developed for conventional and regularized approaches (e.g., Friedman 1989; Hoffbeck and Landgrebe 1996) and are probably extendable to the phylogenetic comparative methods developed here. Also, we introduced penalties for the estimation of the variance-covariance matrix of residuals; adding penalties for the estimation of the coefficients would further improve the statistical performances of the tests and help in the process of predictors selection in (P)GLS analyses. Finally, it would be useful to develop diagnostic tools for detecting outliers in multivariate phylogenetic regression analyses, similarly to what has been proposed for phylogenetic univariate (Revell et al. 2018) and conventional multivariate models (Caroni 1987; Barrett and Ling 1992; Srivastava and von Rosen 1998; Barrett 2003).

With datasets of increasing sizes, further efforts will also be needed to improve computational efficiency; they will be even more needed if regularization techniques with improved performances but more computationally demanding, such as those using quadratic penalties, are required (Engel et al. 2017; Clavel et al. 2019). We already proposed efficient algorithms that can easily be parallelized on modern computing platforms and approximations to the cross-validated log-likelihood for computing the distribution of the statistics. These computations (see Appendix 2) could be further improved by using rank-one update of the eigen-decomposition used in the pre-transformation for the LOOCV (e.g., Mertens et al. 1995). For very high-dimensional datasets (*p*>1000), the computational burden for computing the distribution statistics can be avoided (or reduced) by using PL-based reduction techniques. We previously developed the PL-PCA and were able to apply it on a dataset of *p*=1197 (Clavel et al. 2019). In principle, one could use the ML or PL-MANOVA (with the model estimated when performing the PL-PCA) on a reduced set of these PL-PCA axes. This is a much less computationally demanding test. In this case, the appropriate phylogenetic structure is used in the construction of the pPC axes, and so this procedure is expected to perform well, although we did not directly assess its statistical performances. Another promising direction concerns the probabilistic latent variable models (e.g., Tipping and Bishop 1999; Tolkoff et al. 2018), which could further improve the efficiency of the reduction techniques for super high dimensional comparative data.

## CONCLUSIONS

The development of multivariate phylogenetic comparative tools is urgently needed to study phenomic and more generally quantitative datasets of ever-increasing size across the tree of life. In this paper we identified several limitations of currently available approaches and proposed new tools – GLS based on regularization techniques – that show improved statistical performances in a wide range of situations. We focused here on phylogenetic structure in the residuals, but the proposed approach can be used to handle other forms of correlations in the residuals, such as in time series or spatial data analyses.

## FUNDING

This work was supported by European Research Council grant ERC 616419-PANDA (to HM) and a Marie Curie-Skłodowska Individual Fellowship IF 797373-EVOTOOLS (to JC).

### ACKNOWLEDGEMENTS

The authors wish to thank Leandro Rabello Monteiro for providing the phyllostomid bats dataset. Finally, we thank Leandro Aristide, Carmelo Fruciano, Sophia Lambert, Odile Maliet, Benoît Perez, Ignacio Quintero, Ana C. Afonso Silva, and Guilhem Sommeria-Klein for helpful comments on the manuscript.

## APPENDIX 1: PENALIZED-LIKELIHOOD OPTIMIZATION

The regularized estimator in Equation (6) can be seen as the solution of the following “ridge” penalized log-likelihood (see van Wieringen and Peeters 2016; Clavel et al. 2019):

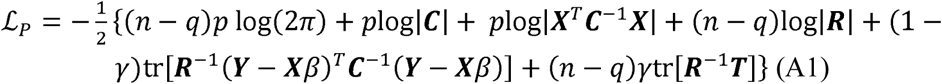

This equation (A1) is similar to the restricted (or residual) log-likelihood described in Clavel et al. (2019; Eq. 1b) but considering the more general case *n − q* rather than *n* − 1 (where *q* is the rank of ***X***) which corresponds to the loss in degrees of freedom resulting from estimating *β* (see details in Harville 1974, 1977; Searle et al. 1992). We find the optimal value for *γ* and the estimator ***R***(*γ*) by maximizing the leave-one-out cross-validated (LOOCV) log-likelihood:

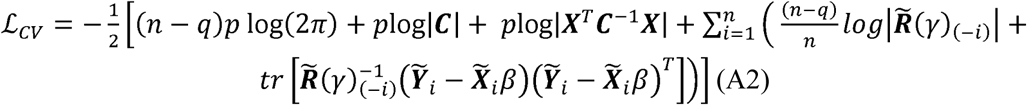

Where 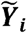 and 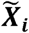 are the *p* by 1 column vectors made of the *i*^*th*^ row of the matrices 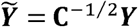 and 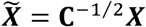 respectively. Thhe *p* by *p* penalized likelihood estimator 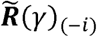 is computed as described in Eq. 6 but with 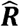 replaced by:

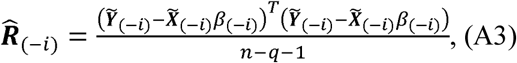

Finding a suitable regularization parameter *γ* is a long-standing problem as it can be achieved in different ways that may optimize different criterion (Ledoit and Wolf 2004; Warton 2008; Hastie et al. 2009). Maximizing the cross-validated likelihood (A2) is known to reduce the statistical risk based on the predictive Kullback-Leibler (K-L) information (Stone 1974, 1977; Yanagihara et al. 2006) and has been shown to perform well with high-dimensional phylogenetic comparative studies by providing accurate estimates of the models parameters (including a well-conditioned estimate of the traits evolutionary covariance matrix ***R***; see details in Clavel et al. 2019).

## APPENDIX 2: EFFICIENT OPTIMIZATION OF THE REGULARIZATION PARAMETER IN PERMUTATED DATASETS

We use derivative-based Newton-Raphson and quasi-Newton algorithms (Nocedal 1980; Byrd et al. 1995) to efficiently find the regularization (or tuning) parameter *γ* that maximizes the cross-validated log-likelihood in Equation (A2) for each of the permuted datasets. Following Warton (2008), we started by expressing the LOOCV log-likelihood (Eq. A2) through the spectral decomposition of the “training” samples covariance matrix 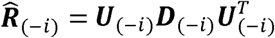 for *i* ∈ [1, *n*] and by using the eigenvector’s matrix ***U***_(*-i*)_ to rotate the residuals of the “test” sample: 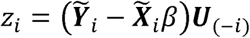. We note that the spectral decomposition of 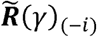 differs from 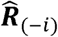 only by a linear transformation of the eigenvalues : 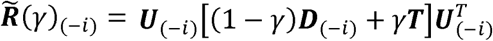 (Warton 2008; Witten and Tibshirani 2009; van Wieringen and Peeters 2016). The parts depending on *γ* in Equation (A2) can thus be expressed as a sum of independent terms that are easier to evaluate:

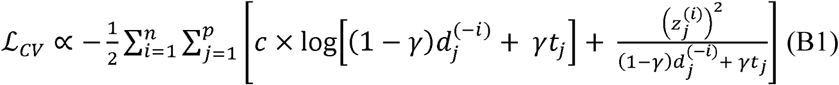

Where 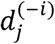 is the *j*^th^ eigenvalue estimated from the eigen-decomposition of the covariance matrix 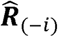, and *c* is a constant equal to 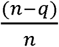 Given the linear form of the cross-validated log-likelihood function (Eq. B1), we can differentiate the summands separately using standard derivative rules to obtain the first and second order derivatives with respect to the regularization parameter *γ* (see Supplementary Material for details). The first derivative is given by:

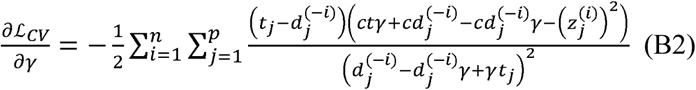

And the second order derivative (used in Newton-Raphson like methods) is given by:

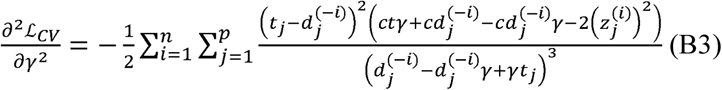

In practice we only used the first derivative (Eq. B2) along with Equation (B1) within a quasi-Newton method (the L-BFGS-B algorithm implemented in the *optim* function in R; Byrd et al. 1995) as it was more efficient than our own implementation of the Newton-Raphson algorithm based on both Equations (B2) and (B3).

Although performing the *n* eigen-decompositions of the LOOCV procedure for the *p* by *p* covariance matrices may appear to be computationally prohibitive, this has to be done only once; repeated evaluations of the cross-validated log-likelihood (Eq. B1), and its derivatives (Eqs. B2-3) during the updating steps of the optimization is done very efficiently given that the dominant computations only involve vectors or scalar operations. It should be noted that this approach assumes that the phylogenetic covariance ***C*** is known and fixed. For instance, with the Pagel’s λ model considered here, we assume that for each of the permuted datasets the phylogenetic covariance ***C*** is known and fixed to the penalized likelihood estimate obtained from the empirical data – that is 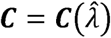. As a result, with the exception of the Brownian motion case, this strategy is generally approximate. We expect the test statistic to be slightly biased (e.g. upwards for the Wilks’s Λ test – McLachlan 1987) since the estimation of the likelihood on each permuted dataset is not based on the maximum likelihood estimate of *C* on this dataset.

